# QUETZAL - an open source C++ template library for coalescence-based environmental demogenetic models inference

**DOI:** 10.1101/214767

**Authors:** Arnaud Becheler, Camille Coron, Stéphane Dupas

## Abstract

The purpose of this article is to introduce an implementation framework enabling us, using available genetic samples, to understand and foresee the behavior of species living in a fragmented and temporally changing environment. To this aim, we first present a model of coalescence which is conditioned to environment, through an explicit modeling of population growth and migration. The parameters of this model can be infered using Approximate Bayesian Computation techniques, which supposes that the considered model can be efficiently simulated. We next present Quetzal, a C++ library composed of reusable generic components and designed to efficiently implement a wide range of coalescence-based environmental demogenetic models.

## 2 Introduction

### 2.1 Motivations

Understanding how species react to spatio-temporal environmental heterogeneity and how this conditions the patterns of genetic variation is of great importance in the context of conservation biology, for example to predict future species distributions under global climate change (Pauls et al., 2013). Spatially explicit simulation studies have proven to be of fundamental importance when tackling such dynamical processes, especially in the context of range expansions (Excoffier et al., 2009). Despite a growing number of simulation programs dedicated to coalescence-based models of genetic variation, code reuse is still limited. We present Quetzal, a new C++ library with reusable generic components designed to ease the implementation of a wide range of coalescence-based environmental demogenetic models, and to embed the simulation in an Approximate Bayesian Computation (ABC) framework. The code is open-source, and available at https://github.com/Becheler/quetzal (see Becheler, 2017).

### 2.2 Context

Present genetic data can be linked to past ecological processes by coupling demographic models accounting for the spatio-temporal landscape heterogeneity with models of genetic variation (see Figure 1). When the studied genetic variation is neutral, genetic models based on coalescent approaches (Nordborg, 2001; Hein et al., 2004; Wakeley, 2009) can be used. In this framework, the coalescence of two gene copies into a parent copy is simply the replication of the ADN viewed backward in time. The genealogy of the sampled genes copies can be defined backward in time conditionally to the demographic process which itself can be defined before tackling genetical aspects. This is an important theoretical link between a genetic sample and the historical processes that shaped it, and it can be used for constructing statistical models allowing to estimate properties of these past processes on the basis of the present sample.

**Figure 1:**
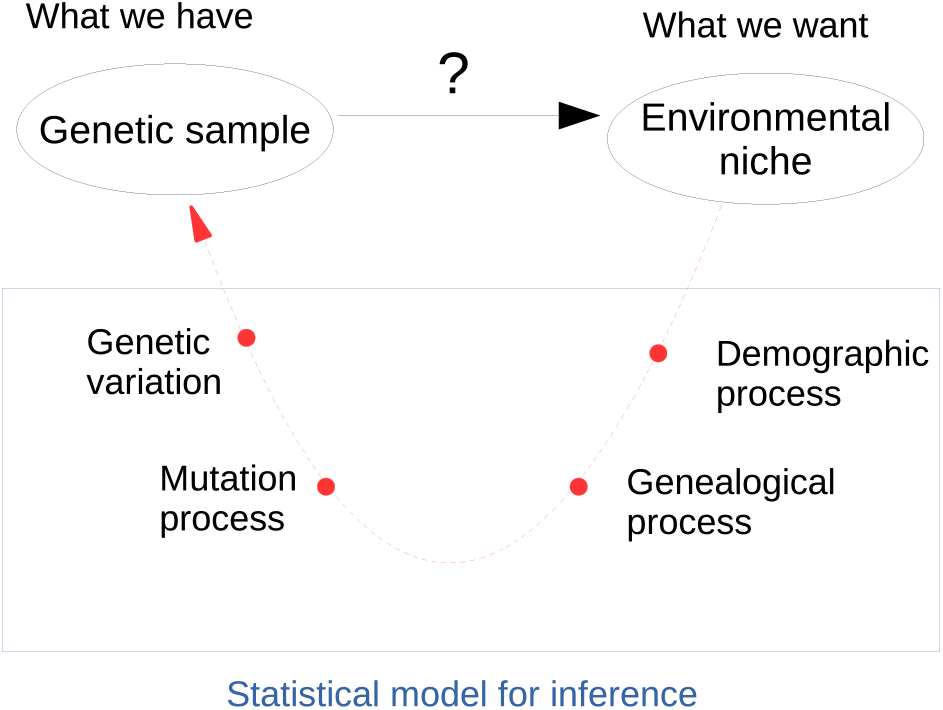
Inferencial framework from which Quetzal stems. The red arrow represents the dependency structure under the hypothesis of neutral variation, where processes condition the observed data. The black arrow represents the inferencial framework, where data allow to shed light on processes.

Constructing such estimates often relies on the study of the likelihood, that gives the probability of data to arise, as a function of the parameter *θ* of the statistical model. The likelihood function can be derived under simple models, but as theoretical advances steered models towards higher levels of complexity (migration (Beerli and Felsenstein, 1999), recombination (Kuhner et al., 2000), selection), the likelihood function became harder and harder to calculate.

Approximate Bayesian Computation (ABC) methods (see Marin et al., 2012) dramatically extended the complexity limits of the models under which inference was possible. ABC bypasses the complex task of evaluating the likelihood function by combining two approximations making the problem computationally tractable: (i) observed data are reduced to lower-dimensional quantities (the so-called summary statistics), (ii) the inference is tolerant to small distortions of the observed summary statistics (see Blum et al. 2013 for more formal explanations). This makes possible for ABC procedures to estimate posterior densities of the parameters by simulating data under the model while exploring the parameter space conditionally to a prior distribution, and accepting only the values of the parameters for which simulated data are close enough to the observations. Despite its apparent ease, ABC methods present important methodological pitfalls (for example the choice of the dimensional reduction function), but many studies have paved the way for the non-statisticians (see Bertorelle et al., 2010) for an excellent methodological guide), and ABC became very popular in Ecology and Evolution (Beaumont, 2010; Csilléry et al., 2010).

The popularity of ABC methods encouraged the development of more complex coalescence-based simulation computer programs, and their authors put tremendous efforts in successfully delivering novative, usefull and user-friendly products to the community of population geneticists. SPLATCHE (Currat et al., 2004) simulates coalescents based on complex demographic simulations run in a spatially explicit landscape, incorporating landscape heterogeneity. Various versions of SPLATCHE largely fostered the rapid expansion of the so-coined iDDC modeling approach (integrated distributional, demographic and coalescent modeling, He et al. 2013a). iDDC uses Approximate Bayesian Computation with spatially explicit demographic simulation model (possibly integrating landscape heterogeneity) to estimate quantities of interest such as populations growth rate or dispersal law parameters (Lacey Knowles and Alvarado-Serrano, 2010; Estoup et al., 2010; Massatti and Knowles, 2016). DIY ABC (Cornuet et al., 2014) is an open-source program that provides the ability to conduct inference under a wide range of complex biological scenarios combining an arbitrary number of admixture, divergence or demographic change events. It offers very strong ABC support and an intuitive Graphic User Interface (GUI). IBDsim (Leblois et al., 2009) is an open-source program for simulating genetic variation under isolation by distance, and provides much flexibility in the choice of dispersal kernels. MSMS (Ewing and Hermisson, 2010) puts emphasis on incorporating selection and proposing extensible design. These programs, and others, have provided invaluable support to the non-developer communities for a wide range of applications and studies.

Paradoxically, these programs collectively failed to help the software developers community to write new programs, mainly due to very low rate of reusability in their source code. Because they put emphasis on the non-developer user, they act as rigid black boxes taking an input, processing it in some way configured by some form of User Interface (*e.g.* command line or GUI) and delivering the output. Indeed they can not be used if the underlying theoretical model does not belong to their predefined set of possible options (*e.g.* SPLATCHE does not allow to change the dispersal kernel or the local growth model) or if their computational solution does not answer to the question (*e.g.* if they write standard genetic summary statistics in output files when we would need to analyze genealogical properties). This current state of the art does not scale with the virtually infinite number of arbitrarily complex evolutionary or demographic models and the evergrowing number of statistical methods variants. We need standard, general, reusable tools for helping us to quickly build programs that can solve new problems. As written in Stroustrup (2003): “The key to fast development, correctness, efficiency, and maintainability is to use a suitable level of abstraction supported by good libraries“. Reusable code such as library’s relies on abstractions, syntactic constructions often opposed to performance. However, when it comes to ABC and massive simulations, performances become critical. C++ offers the template mechanism, a key feature allowing to build very high levels of abstraction without loss of efficiency, so we do not have to choose between reusability and performance.

As a first attempt to offer reusable components to the coalescence software community, we present here Quetzal, an open-source C++ template library. Quetzal offers powerful abstractions for building coalescence-based simulation programs. It contains several independent modules, each directed towards a general simulation purpose (demography, geography, coalescence, genetics, ABC …) in which are located the files containing the sources of generic components. The extensive documentation and the wiki (Becheler, 2017), both avalaible on the github project, ease to pick and combine the desired functionalities and the template mechanism allows to adapt it to a new problem with minimal recoding (if no recoding at all). We insist on the possibility to extend the behaviors of the Quetzal algorithms with very few new lines of code. The high level of abstraction in Quetzal allows its generic components to be used to write expressive and maintainable code with appreciable terseness and efficiency. Their genericity make them applicable to a large range of programs and the user can change the design of its application without major changes in the code (what typically arises when changing a finally-not-so-minor detail in the theoretical model). Each documented functionality comes with a small demonstration program and its output, providing valuable and intuitive insights on the way to manipulate the component. The library design insists heavily on modularity, extensibility and efficiency, and is intended to respect the standards of the Standard Template Library with STL-like algorithms and interfaces. Template rules make the library header-only.

This first version focuses on coalescence-based environmental demogenetic models. Consequently, Quetzal first features are designed to bring efficiency and flexibility in defining demographic quantities as functions of space and time of landscape heterogeneity, to couple these quantities to the coalescence process, and to use ABC methodology to conduct inference.

First we present a simple mathematical ecological model for estimating ecological features of a species from a genetic sample with the ABC procedure. This model is purposely general, as it will serve as an illustration to facilitate the understanding of Quetzal functionalities by enforcing its genericity, but it is still of strong biological interest as it relaxes several constraints that were previously made in the literature. Of course Quetzal is flexible and its use is not limited to this model as other features can be added or removed. Then we present a core feature of Quetzal for the coalescence, the abstraction of the ancestry relationship between a child gene copy and its parent, and we point out its importance to use and extend the library. Lastly, we give an overview of Quetzal functionalities and concepts and we apply them in a demonstration program implementing a fully-specified version of the previous general theoretical model.

## 3 Ecological and mathematical demogenetic model

### 3.1 Motivations

Let be a spatial sample *S* of *n* haploid individuals that have been sampled at time *t*_*s*_ across the landscape and that have been genotyped at a microsatellite locus, and a dataset giving the values of environmental quantities across the same landscape. From these data, we want to infer ecological properties of the species such as the *niche functions* (defined here as the functions relating the environmental quantities to demographic quantities, for example the local growth rate) and the dispersal function, so we need a statistical model to link observed data and processes to infer (see Figure 1). We present here a description of a general bayesian model for estimating these functions using an ABC framework (see Figure 2): a demographic history is simulated forward in time conditionally to the features of the heterogeneous landscape, then the genealogy of the sampled gene copies is simulated backward in time conditionally to the demography. The uncertainty on the parameters 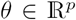 to estimate is defined by the prior distribution II from which a parameter *θ* is sampled at each simulation.

**Figure 2:**
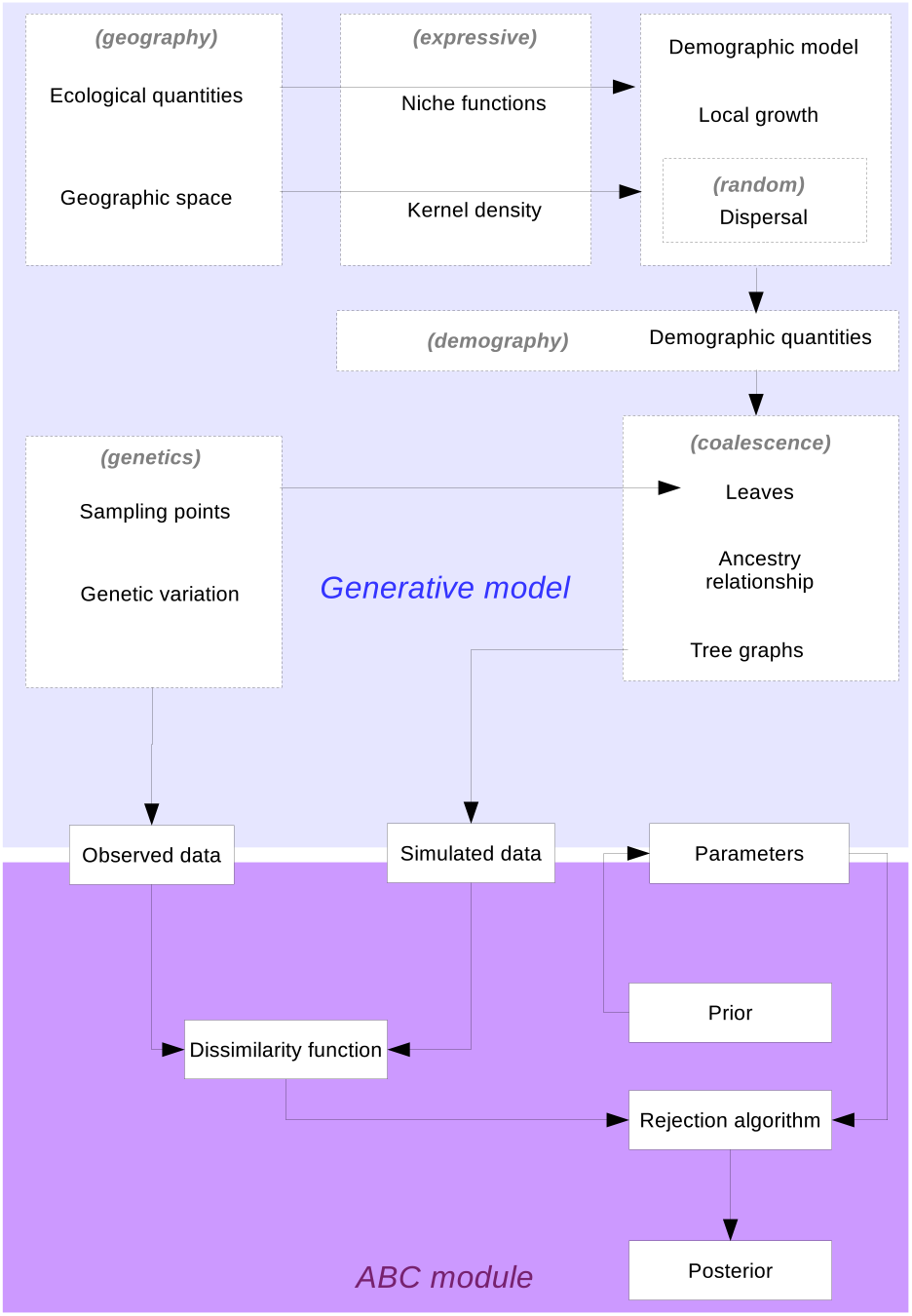
Flow chart illustrating the information flows between components of a general environmental demogenetic model (see section 3). In grey parenthesis are indicated the Quetzal modules (see section 5) that allow to represent these components in the code. The way the landscape conditions the demographic processes form the main focus of a number of approaches (landscape-ABC (Estoup et al., 2010), iDDC modeling (He et al., 2013b)) in the literature, such that the inference is usually driven on the underlying niche and/or dispersal model. Infering such ecological properties from a spatial genetic sample is made possible by using a coalescence model to link the sample to the demographic processes that shaped it. Inference is run in an ABC framework, where parameters to estimate are sampled in a distribution, allowing a dataset simulated by the generative model to be compared to the observed data by some dissimilarity function to build the posterior.

### 3.2 Geography

Let consider a given set of demes 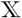 (typically reduced to the geographic coordinates of their centroïd). The environment *E* is defined by *i* known ecological quantities which are functions of space and time, typically climate layers from the WorldClim global climate database (www.worldclim.org), or a niche suitability dataset estimated from an external niche modeling step (He et al., 2013a).

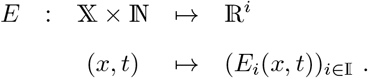

### 3.3 Demography

The demographic simulation process goes from time *t*_0_ to *t*_*S*_ and iteratively constructs the function *N* giving the number of individuals in deme 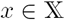 at time *t:*

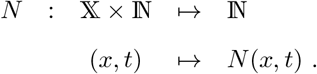

*N* is initialized by setting *N*(.,*t*_0_) the initial distribution of individuals across demes at the first time *t*_0_-Typically for a biological invasion, this is restricted to the introduction site(s) with the number of introduced individuals (Estoup et al., 2010). For endemic species, paleoclimatic distribution can be considered as starting point. The number of descendants 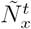 in each deme is sampled in a distribution conditionally to a function of the the local density of parents, for example 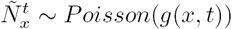, where *g* can be for example a discrete version of the logistic growth as in Currat et al. (2004).

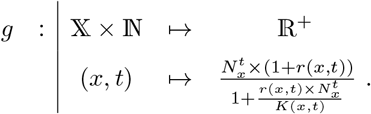

The *r* (respectively *k*) term is the growth rate (respectively the carrying capacity), defined as a function of the environmental quantities with parameter *θ:*

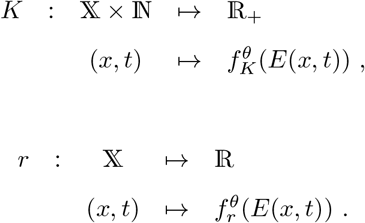

Non-overlapping generations are considered (the parents die just after reproduction). The children dispersal is done by sampling their destination in a multinomial law, that defines 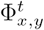 the number of individuals going from *x* to *y* at time *t:*

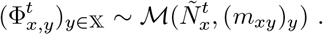

The term (*m*_*xy*_)_*y*_ denotes the parameters of the multinomial law, giving for an individual in *x* its proability to go to *y.* These probabilities are given by the dispersal law with parameter *θ:*

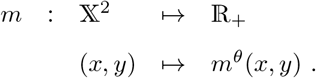

After migration, the number of individuals in deme *x* is defined by the total number of individuals converging to *x:*

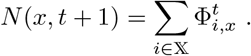

### 3.4 Coalescence

These quantities are used for defining the coalescence process which is defined by the following stochastic process going from *t*_*s*_ to *t*_0_: knowing that a child node *c* is found in 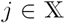 the probability for its parent *p* to be in 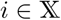 is :

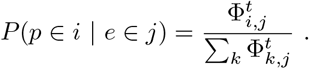

Knowing that the parents *p*_1_ (*p*_2_) of nodes *c*_1_ (*c*_2_) are in *x* at time *t*, the probability for the children to coalesce in the same parent is :

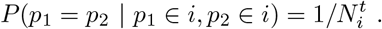

A forest of random coalescent trees is then constructed backward in time, until *t*_0_ is reached. Note that at this point the Most Recent Common Ancestor is not necessarily found, and assuming that *t*_*S*_ − *t*_0_ is small enough no neglect mutations, the simulation can end with a collection of trees rather than a complete genealogy.

## 4 Abstraction of the ancestry relationship

### 4.1 Motivations

When exposing the concept of coalescent trees above in the mathematical model, it has been useless to define exhaustively the tree properties or the exact nature of its nodes and branches. These are details humans typically *abstract away,* which leads to high generalization and low intellectual overhead. However, a computer program has to deal with an impressive number of details, and if these details are not carefully separated from the general concerns when writing the code, it leads to poor generalization (see Alexandrescu, 2001, p.xvii). Indicators of a lack of generalization are typically numerous dependencies across code, monolithic classes and a high rate of code duplication. Consequences are defined by Martin (2000) and Alexandrescu (see 2001, p.5) as being *rigidity* (the software is difficult to change), *fragility* (the software breaks at several points after a small change), *immobility* (the software can not be reused in another context, so it is entirely rewritten), gruesome intellectual overhead and poor performances. Since the devil is in the details, a natural solution is to write code in terms of general *abstractions* rather than in terms of *implementation details* (Dependency Inversion Principle, Martin 2002), so the designed generic components can be reused in various contexts.

### 4.2 Object-oriented paradigm

In C++, the genericity of the implementation can be realized by using inheritance and dynamic binding enable by the virtual keyword (Object Oriented Programming, OOP). Algorithms manipulating trees would then rely on an interface defined in terms of an abstract class `AbstractTree`, but will be applied on instances of concrete classes that inherit from `AbstractTree` and that present the specific desired behaviors, for example `TreeForStoringCoalescenceTimes` or `TreeForStoringCoalescenceDemes`. This design avoids the well-known problems of a class that would expose a monolithic interface to store all possible features like demes, times, mutations and others, (Single Reponsability Principle, Martin 2002) but it has a number of well-known drawbacks. First, if inheritance can be very useful when the set of classes to be treated by the algorithm can naturally be thought as a hierarchy of concepts, in most cases this is not the case: it would result very unnatural to order them into a class hierarchy, and, importantly, it would lead to hardly maintainable code design (Stroustrup, 2014). Second, it is sometimes natural to expect the algorithms to work with primitive types or with STL containers, but as primitive types are not classes and as STL containers are not designed for dynamic polymorphism (they have no virtual destructor), there is no hope to see the object-oriented approach work with these types. Finally, the use of inheritance and virtual functions can have runtime overhead because of an extra lookup in the virtual table when a virtual function is called, and because virtual methods can not be inlined (Stroustrup, 2014).

### 4.3 Generic paradigm

All the data type manipulated by a same general algorithm do not have to be linked by the rigid hierarchical relation imposed by OOP : the generic programming allows for uniform manipulation of independent types. In generic programming an algorithm is not defined in terms a particular type, but in terms of a set of constraints wielded on the type by the algorithm internals; this set is frequently defined as a *concept* and represents an implicit interface. Thus the algorithm will work with any type fulfilling these constraints, allowing for high abstraction level without loss of efficiency. Generic programming allows to implement the coalescence algorithms with great generalization, making minimal assumptions on the type handled.

The simple task to merge two nodes uniformly at random in a sequence of nodes can be defined as follow:

1. randomly permute the *k* elements of the sequence
2. create of a new node *P* (the parent)
3. designate the first element as child of *P*
4. designate the last element as child of *P*
5. assign *P* to the first element.
6. return the *k* - 1 first elements.

When implementing this binary merge algorithm in Quetzal, several details need to be abstracted away to preserve the generality of the algorithm definition: the nature of the nodes, the nature of the sequence, of the nature of the inheritance relationship between a child node and its parent.

The nodes could be integers, character strings, a user-defined class or, actually, *anything else*. No constraint on this type comes from the algorithm, but various constraints can come from the sequence type used to store them. The sequence could be a standard container (`std::vector, std::list` …) or a user-defined type. The classical way to abstract containers in C++ is passing as argument two iterators (one pointing to the first element of the sequence and the other pointing to the past-the-end element) giving the range of data on which the algorithm will operate. As the algorithm can not modify the external container, the reduction in size caused by the merge is signaled by returning an iterator pointing to the new past-the-end element. The only explicit effort the user can have to do is just to precise, conditionally to the chosen node type, what is meant by “*designating node c as child of the parent node p*“. This can done by passing to the algorithm a function-object taking as argument a reference on the parent and another on the child, returning the result of the branching event. Or, if no function-object is given, the sum operand is used by default (thus requiring the expression *c* + *p* to be defined).

This abstraction of the ancestry relationship is expected to allow (i) to efficiently generalize the existing Quetzal algorithms to an open-ended number of specific, user-defined kinds of genealogies and (ii) to give guidance to the developers who need to write new generic coalescence algorithms.

#### 4.3.1 Count hanging subtrees leaves

Achieving genericity is of fundamental importance to efficiently tackle a wide range of situations. For example most of the current simulation softwares focuse on generating genetic variation samples, because this is most of the time the only information available. However, in some cases the mutational process can be negligible compared to the recent genealogical process in shaping the sample configuration (see Becheler et al., in preparation), so topological properties like the number of leaves of hanging subtrees (see Hein et al., 2004, p.78) become the desired output. This is very unlikely that the developer of a program could ever foresee this specific need. Fortunately, it does not mean one should recode everything from scratch each time a new simulation behavior is needed by a new methodological advance: Quetzal allows the user to inject the desired behaviors into its generic components.

When implementing the simulation, instead of building complex genealogical objects, then counting their leaves by tree traversals algorithms, a much more efficient approach is to directly make the coalescence algorithm sum the number of leaves of the hanging subtrees, updating it at each coalescence event. The type of nodes is thus defined as being integers, and the sampled nodes value is set to 1. Conveniently and by default, the merging algorithm will initialize the new parents to their default constructor value (which is 0 for integers), and define the branching event of two nodes by the sum function.

The following small program applies the approach by merging two nodes uniformly at random in a sequence of four nodes, updating the leaves number information. It can of course be extended to much more complex simulation frameworks.

~~~
#include "quetzal/coalescence.h"
#include <random> // std::mt19937
#include <iostream> // std::cout
#include <algorithm> // std::copy
#include <iterator>
using namespace quetzal::coalescence;
int main(){
 using node_type = int;
 std::vector<node_type> nodes(4,1);
 std::mt19937 rng;
 auto last = binary_merge(nodes.begin(), nodes.end(), rng);
 std::ostream_iterator<node_type> it ( std ::cout, "u");
 std::copy(nodes.begin(), last, it);
 return 0;
}
~~~

The output gives the number of leaves of each hanging subtree after one generation of coalescence: 2 1 1

#### 4.3.2 Construct a Newick tree format

Code with a suitable level of abstraction allows to readapt old code to new problems with ease. Studying genealogies topological properties can be the main statistical focus, but visualizing genealogies is still the most instinctive way to shed light on some properties, to assert correctness of algorithms generating them, or to present results. However, when it comes to data visualization, C++ is not the most suited platform. Many tree visualizer use a Newick tree format (see Olsen, 1990) as an input for nice plot rendering. The implementation is straightforward when using Quetzal abstractions: the type of the nodes is now a character string, the parent node is by default constructed as an empty string, and the branching event is defined as a formating function taking the parent node *p* and the child node *c* as argument to build the Newick format character string piece by piece. There are very few lines to change in the code to entirely redefine the meaning of a coalescence event:

~~~
#include "quetzal/coalescence.h"
#include <random> // std::mt19937
#include <iostream > // std::cout
#include <algorithm > // std::copy
#include <iterator >
#include <string >
using namespace quetzal::coalescence;
int main(){
 using node_type = std::string;
 std::vector <node_type > nodes = {"a","b","c","d"};
 std::mt19937 rng;
 auto branch = [](auto p, auto c){
  if(p.size() == 0)
  return "(" + c;
 else
  return p + "," + c + ")";
 } ;
 auto first = nodes.begin ();
 auto last = nodes.end();
 while(distance(first,last)>1){
  last = binary_merge(first, last, rng, branch);
 }
 std::ostream_iterator<node_type> it ( std ::cout, "u");
 std::copy(nodes.begin(), last, it);
 return 0;
}
~~~

The output will give the Newick format character string ((d,a),b,c) representing the following coalescent tree.

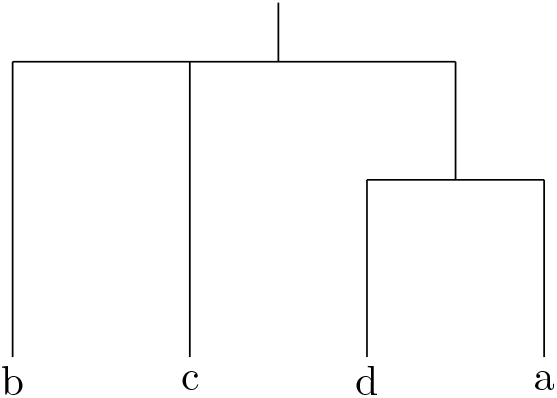

The output can be exported for example on an online tree viewer such as iTol (Letunic and Bork, 2006). Quetzal is not restricted to this simple example and any arbitrarily more complex formatting functions can be considered, for example to represent branches length or nodes position in a landscape.

## 5 Quetzal components for simulation

The manipulation of the genealogies is the most fundamental aspect of all coalescence-based application programs, so the abstraction of the ancestry relationship is expected to be always useful, and the algorithms written in terms of this abstraction are expected to be highly reusable. However, an open-ended number of generative model variants can be considered: we present here a number of components that are most likely to be necessary when implementing them. Note that these components are intended to be independent. For example, if a demographic simulation is usually run after reading some environmental quantities in a geographic file, this does not have to be the case. Indeed any user-defined set of coordinates can be used to represent the demic structure, and environmental quantities can for example be represented by any mathematical function of the geographic space. Accordingly, the type of the geographic coordinates used in the *geography* module does not pervade the other modules.

### 5.1 Discrete landscape construction

In the *geography* module, Quetzal uses the Geospatial Data Abstraction Library (GDAL Development Team, 2017) to read grids of ecological data through the instantiation of a `DiscreteLandscape` object. In this class, the demes are represented by the grid cells and identified by the geographic coordinate of their centroid. The class allows to retrieve the set of demes centroid geographic coordinates (so it can be used to represent the demic structure to run spatially explicit simulations), to reproject a set of sampled coordinates to the nearest centroid (so compatibility is ensured between a spatial sample and the geographic support), or to deliver lightweight function-objects that give access to the underlying ecological quantities and that are susceptible to be coupled to the demographic model by composing them into arbitrarily complex mathematical expressions of space and time. The `GeographicCoordinates` class allows secured manipulations of longitude and latitude coordinates, and computation of great circle distances (often useful to define dispersal kernels).

### 5.2 Demographic variables definition

In the *demography* module, Quetzal defines two class templates (`PopulationSize` and `PopulationFlux`) to construct and consult 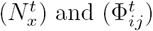, providing expressive interface for secured and intuitive manipulation. Both class templates do not depend on *geography* module as they are templated on the type of the locations (the demes) and on the type of values that are stored. It makes possible to use any geographic coordinate system (for example longitude/latitude using the `geography::GeographicCoordinates`) to spatialize the model. Any arithmetic type can be chosen to define the set in which 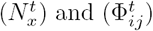 take value (typically 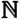 or 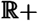). Indeed, the type of the stored time values is not necessarily an integer but can be a more complex date type.

### 5.3 Compile-time functions composition

Because growth patterns are species-specific features, the expression of 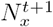 is typically user-defined and hence can be any arbitrarily complex function, for example constant values or discrete version of logistic growth model (Currat et al., 2004). Quetzal offers tremendous facilities to build these functions, composing function-objects into an expression that can be efficiently passed around. To this purpose, Quetzal integrates *expressive* (Marques, 2017), a library making use of metaprogramming techniques to enable compile-time optimization of function composition with high expressiveness.

Consider the discrete logistic growth version (see section 3) to define 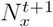 and let us pretend there is strong biological motivations to want the growth rate *r* to be constant (*r* = 3) over space and time, and the carrying capacity *K* to be the mean of heterogeneous environmental quantities 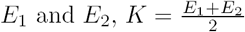. As C++ has strong static typing, it is impossible to directly sum constants (literals) with functions, as it would be possible under others languages. So the first step is to transform the constants as functions with the same definition space as the other functions we want to combine:

~~~
literal_factory <space_type, time_type > lit;
auto r = lit(3);
~~~

Here *r* is now callable with time and space arguments, and can be composed with other *expressive* callable objects. Assuming that *E*_1_ and *E*_2_ are function-objects callable with time and space arguments (for example the function-objects delivered by the `DiscreteLandscape` class), the function use allows *expressive* to manipulate the expressions and gives them an enriched mathematical interface, so the addition or division operator can be applied on them:

~~~
auto K = (use(E_1)+use(E_2))/lit(2);
~~~

And finally, assuming that *N* is a function-object callable with space and time arguments, the whole growth expression can be built:

~~~
auto g = (use(N)*(lit(1)+r)/(lit(1)+((r * use(N))/K));
~~~

This expression can be captured in a lambda expression to simulate the number of gene copies after reproduction in deme *x* at time *t*:

~~~
auto sim_growth = [g](auto x, auto t, auto& gen){
  poisson_distribution <N_type > dis(g(x,t));
  return dis(gen);
}
~~~

This object can be passed around conveniently for further invocation with space and time arguments:

~~~
for(auto t : times){
 for(auto x : space){
  // …
  auto N_tilde=sim_growth(x,t,gen);
  // …
 }
}
~~~

This last code snippet illustrates an important benefit of *expressive*: to allow the separation of concerns when writing the application code. In other terms, the details of the logistic growth expression are not intricate with the code where the expression is invoked, what would result in an obfuscated and hardly maintenable code. Moreover, as the expression is known at compile-time, it is a perfect candidate for all compile-time optimizations done by the compiler such as inlining: as the compiler knows exactly which functions will be called when *g* is called, he should be able to replace a function call directly by the body of the function, potentially leading to extremely efficient code with very few indirections.

### 5.4 Dispersal patterns

In the *random* module, the `TransitionKernel` class is an implementation of a markovian kernel for sampling the next state *x*_1_ of a markovian process conditionally to the present state *x*_0_. It can be used in the dispersal context (in that case the states type will represent demes coordinates, for example `geography::GeographicCoordinates`). The underlying markovian probability distributions associated with each present state do not need to have the same arrival space, and they are built only if needed by the simulation context. The weights are computed with an arbitrary mathematical function conditionally to the present state (for example when only geographic distance affects dispersal) or to the present state and time (for example when environmental spatio-temporal heterogeneity affects dispersal).

### 5.5 Coalescence features

For coalescence under the Wright-Fisher model, a binary merge algorithm is proposed to be used in the simulation contexts where the sample size is small relative to the population size. A simultaneous multiple merge algorithm can be used when this approximation does not hold.

The benefit of abstracting the inheritance relationship when simulating the genealogical graph was presented above along with two examples showing that an explicit representation of the coalescent was not necessarily desirable. However in many standard cases, it is needed, for example to save arbitrary information from the simulation context (times of coalescence events) and access them later for updating some quantities while descending the genealogy (for example apposing mutations with a probability conditional to the time spent between two nodes). For these cases, the Tree class template allows to construct such object, encapsulating an arbitrary user-defined data field into each node, defining the inheritance relationship between a parent node and a child node in a secured way, proposing topological manipulation operations and tree traversal algorithms.

The Forest class template is designed to ease the manipulation of spatial collections of trees (of arbitrary type) when using spatially explicit coalescence simulation.

## 6 Quetzal components for inference

The Quetzal *abc* module provides abstractions allowing to embed efficiently an open-ended range of simulation models into an ABC framework. To this purpose, an ABC object associates a simulation model and the prior distribution of its (possibly multidimensional) parameter to conduct inference. We present here key elements of the *abc* module, notably the concept of *GenerativeModel* used to abstract out the model-specific details.

As ABC-based inference on spatial coalescents involves complex functions for dimensional reduction and distance computation that are far beyond the scope of this article, the ABC inference examples will be presented with a toy generative model (the poisson distribution), a toy dimensional reduction function *η* (identity) and a toy distance *ρ* (absolute value of the difference).

Then we step away from the toy-model and propose a concrete example of a class satisfying *Generative-Model* and implementing a fully-specified version of the general theoretical model of coalescence presented in section 3. This example will make use of Quetzal components to illustrate how to build original coalescence simulations objects with ABC-compatible interface.

### 6.1 Features

#### 6.1.1 The *Generative Model* concept

To achieve genericity and propose clear, standard and uniform ways to manipulate models and parameters, specific simulation models are abstracted to the concept of *GenerativeModel*, a Quetzal C++ concept that has been designed to be a generalization of the standard C++ *RandomNumberDistribution*. It notably generalizes the type of the result that is no more restricted to be an arithmetic type, so more complex type values (coalescents, genetic data, or summary statistics) can be generated. Furthermore, the returned values are not necessarily generated from a simple probability density function or a discrete probability distribution, as generally in ABC a complex stochastic simulation function is involved. The list of all requirements can be found in the documentation. Any model object whose type D satisfies *GenerativeModel* and any prior object able to randomly produce an object of type `D::param_type` can be used to build an `ABC` object. Consequently, all the STL random number distributions are compatible with the *abc* module, which is very convenient for testing and demonstration purposes:

~~~
using model_type = poisson_distribution <>;
uniform_real_distribution <double > prior (1.,100.);
model_type model;
auto abc = make_ABC(model, prior);
~~~

Here an ABC object is constructed by associating the STL poisson distribution with a prior on its parameter, the STL uniform distribution, for sampling parameters in [0,100].

#### 6.1.2 Prior predictive distribution sampling

The generation of the reference table is done by sampling *n* results in the prior predictive distribution.

~~~
mt19937 g;
auto n = 1000000;
auto table = abc.sample_prior_predictive_distribution(n,g);
~~~

The generated `ReferenceTable` object can compute other table objects. Considering a function-object representing the summary statistics function *η*, the raw data table can produce a second `ReferenceTable` object associating the parameter value to generated summary statistics.

~~~
auto eta = [](auto x){return x;};
auto sumstats = table.compute(eta);
~~~

Generated data can be accessed, for example to be used as pseudo-observed data in ABC validation methodology:

~~~
auto pod_value = sumstats.begin()->value();
auto pod_param = sumstats.begin()->param();
~~~

Then, considering a function object representing *ρ*, the distance function between observed and simulated dataset,

~~~
auto rho = [](auto obs, auto sim){return abs(obs - sim);};
auto distances = sumstats.compute_distance_to(pod, rho);
~~~

Finally the syntax of the various interfaces make it intuitive to design a quick rejection algorithm, sending to output only the parameter values for which the generated summary statistics was less than a threshold:

~~~
double threshold = 2.0;
for(auto const&it : distances){
 if(it.value() <= threshold){
  cout << it.param().lambda() << endl;
 }
}
~~~

#### 6.1.3 Rejection samplers

The simplest samplers is the Rubin rejection sampler (Rubin, 1984). It accepts a parameter value only if the generated data is strictly equal to the observed data (the data type has to be *EqualityComparable*, *i.e.* having a built-in or a user-defined comparison operator `operator==`).

The implementation of the Pritchard rejection sampler (Pritchard et al., 1999) generalizes the dimensional reduction function (that traditionally computes summary statistics) and the distance function (that evaluates the distance between observation and simulation). Therefore, any object-function can be used to transform data into summary statistics and any type of distance can be used.

More complex sampling algorithms like MCMC-ABC (Marjoram et al., 2003), SMC-ABC (Del Moral et al., 2006), or PMC-ABC (Beaumont et al., 2009) are yet not implemented, but we do not expect that it will hinder Quetzal reliability. Indeed these algorithms are known to be challenging to calibrate (Marin et al., 2012), while embedding the simulation model and the inference framework in the same C++ application code has the benefit to make all the type information available for the compiler, decreasing the computational cost, so the generation of the reference table alone is expected to be useful for a wide range of situations. Furthermore, we expect that if more sophisticated versions of algorithms are needed, the Quetzal existing concepts will greatly ease their implementation.

### 6.2 Implementing a custom generative model

We present here how to construct a class `ExampleModel` that meets the requirements of the *GenerativeModel*. The main general ideas are highlighted here and the program can be found in the supplementary material.

We consider a landscape reduced to two demes *A* and *B*. At time *t*_0_, *N*_0_ = 10 haploid individuals were introduced in deme *A*. The local growth rate and the local carrying capacity *K* are assumed to be constant across the landscape. The growth rate is known (*r* = 100) while the local carrying capacity *K* is unknown and assumed to belong to [1, 500]. The aim of the program is to estimate this value starting with a uniform prior distribution on [1, 500]. For each individual there is a probability *m* = 0.1 to migrate to the other deme. After *g* = 10 generations, *n* = 30 individuals were sampled in *B* and genotyped at one locus. We assume that each introduced individual had different allelic states and that mutational process is negligible. Under these hypotheses, the observed clustering of the data is only shaped by the genealogical process, so we can reject all simulated coalescent forests that do not clusterize the dataset into as many subsets of same cardinality than in the observed clustering. Consequently, we just need to construct the vector of the hanging subtrees leaves count (a way to do it efficiently was presented above) and to compare it to the observed vector of clusters size. We accept the parameter used for the simulation only if the two vectors are equals. For a demonstration purpose, we construct a reference table by sampling 5×10^5^ simulated data into the prior predictive distribution, and we generate 100 pseudo-observed data under the parameter *K* = 50 for validation. Note here that if the pseudo-observed data do not contain 30 individuals in deme *B*, no posterior will be estimated. The prior and posterior distributions are shown in Figure 3.

**Figure 3:**
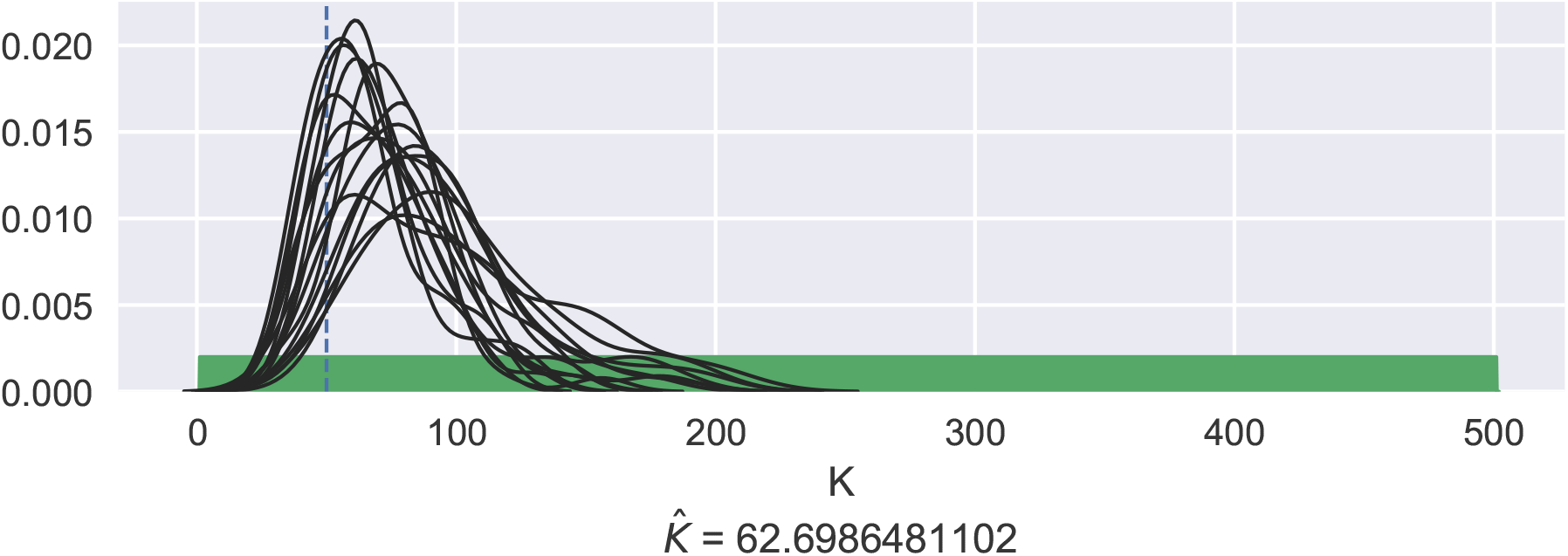
Posterior densities obtained by ABC inference conducted on pseudo-observed data generated under the *ExampleModel* model shown in section 6.2. True parameter value *K =* 50 is shown as the vertical dashed line. The prior distribution is shown in green. The mean of all posteriors is given as 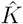.

#### 6.2.1 ABC-compatible interface

Declaring the following interface is sufficient to capt the generality of all possible generative models and enable the `ExampleModel` class to interact with an `ABC` object:

~~~
class ExampleModel{
public:
 using param_type = Param ;
 using value_type = … ;
 template <typename Generator >
 value_type operator()( Generator& gen, param_type const& p) const;
};
~~~

The member type `value_type` describes the type of value generated by the model. The member type `param_type` encapsulates the details of *θ*, the multidimensional parameter to estimate.

#### 6.2.2 Encapsulating *θ*

The point here is to hide useless details (such as dimensionality) from ABC procedures while allowing the user to have control over its implementation. We warn against using vectors or arrays to represent multidimensional parameters: it would impose all dimensions to be of same type, and their index-based value access interface would later favor confusion between dimensions. Instead, we suggest to follow the STL standards and to implement user-defined small classes, with expressive getter/setter syntax. Here we show an extract of the `Param` class, where the member `k` represents *K*, the carrying capacity of each deme in the landscape. Other dimensions are not given here, but can be set to constants for better code maintainability, as found in the supplementary material.

~~~
class Param{
private:
 double K;
public:
 double K() const; // getter
 void K(double); // setter
 // … other dimensions
};
~~~

#### 6.2.3 Constructing the prior

Instances of this parameter class can be created in a prior distribution *i.e* a function that can be called with a random generator and that randomly produces a `param_type` object, manipulating the `Param` object *via* its interface to set its dimension values:

~~~
auto prior = [](auto& gen){
 ExampleModel::param_type params;
 params.k(std::uniform_int_distribution <>(1,500)(gen));
 params.r(100);
 params.m(0.1);
 return params;
};
~~~

More guidance in the design of this second-order function can be found in the project wiki. This function-object, representing the parameter joint distribution, will be passed to the `ABC` object, that will use it to generate random parameters and pass them to the model `ExampleModel::operator()` member function to generate random `value_type` objects. The model details lay in the definition of `ExampleModel::operator()` member function, and a possible implementation is proposed in the supplementary material.

## 7 Acknowledgements

We thank Ambre Marques who importantly contributed to the present Quetzal state by taking on her free-time to provide advice on generic paradigm and design issues, and to develop the *expressive* library.

We thank Florence Jornod who participated as an intern.

Arnaud Becheler was funded by a multidisciplinary project founded by the French Government (LabEx BASC, ANR-11-LABX-0034) that aims to provide new knowledge regarding the drivers of species distribution and to design innovative guidelines toward sustainable land management.

This work was partially supported by the Chair "Modélisation Mathématique et Biodiversité" of VEOLIA-Ecole Polytechnique-MNHN-F.X., by the Mission for Interdisciplinarity at CNRS and by the Institute for the Diversity, Ecology and Evolution of the Living World.

## 8 Data Accessibility

Quetzal source code can be found on github project (https://github.com/Becheler/quetzal). The README file redirects towards Quetzal resources (documentation, wiki, IRC channel). This program is a free software; you can redistribute it and/or modify it under the terms of the GNU General Public License as published by the Free Software Foundation; either version 2 of the License, or (at your option) any later version.

## 9 Authors Contribution

All authors participated equally in the mathematical model design. Arnaud Becheler implemented the C++ library. The article and the documentation of the Quetzal project were written by Arnaud Becheler in cooperation with the other authors.

